# Insulin exerts epigenetic control of joint-specific memory T cells in rheumatoid arthritis

**DOI:** 10.1101/2025.04.01.646580

**Authors:** Venkataragavan Chandrasekaran, Malin Erlandsson, David Svensson, Eric Malmhäll-Bah, Sofia Töyrä Silfverswärd, Gergely Katona, Rille Pullerits, Maria I. Bokarewa

**Author notes:** **Corresponding author:** Maria I. Bokarewa, Department of Rheumatology and Inflammation Research, Institute of Medicine, University of Gothenburg, Box 480, 40530 Gothenburg, Sweden.

## Abstract

**Background:** Insulin has epigenetic effect influencing gene expression. High peripheral insulin concentrations promote insulin resistance in autoimmunity. Oncoprotein survivin/*BIRC5* modulates glucose metabolism through chromatin binding and propagates IFNg effects in CD4^+^ cells. In this study, we explored how insulin influences chromatin binding and metabolic activity in autoimmune CD4^+^ cells of patients with rheumatoid arthritis (RA).

**Methods:** We profiled the metabolic activity of CD4^+^ cell clusters using single-cell transcriptome analysis in blood, synovial fluid and synovial tissue of RA patients. Through chromatin immunoprecipitation and sequencing, we identified the genes controlled by deposition of survivin and acetylated lysine 27 on histone H3 (H3K27ac) in CD4^+^ cells. Treating CD4^+^ cells with insulin and histone deacetylase inhibitors (HDACi), we identified changes in H3K27ac, linked those to transcription of the H3K27-survivin-controlled genes and the pathogenic phenotype of CD4^+^ cells using flow cytometry. Finally, we explored if anti-diabetic and anti-rheumatic drugs affect the metabolic profile and memory phenotype of the metabolic active CD4^+^ cells.

**Results:** Transcription of survivin/*BIRC5* and histone acetylation enzymes strongly correlate with active metabolism in blood CD4^+^ cells of RA patients. In RA synovial tissue, these *BIRC5^hi^* active T cell clusters are inflammatory, exhausted, and memory-like. Genome co-deposition of H3K27ac-survivin pinpointed the insulin-dependent genes in metabolic active CD4^+^ cells. These genes favored histone acetylation by suppressing methylating enzymes *EZH2* and *KMT2A*, and T cell development by activating *CD27, CD3G, and SCIMP*. Inhibition of histone deacetylation reverted these transcriptional effects and supported cellular sensitivity to insulin. Insulin stimulation increased H3K27ac and together with HDACi, suppressed *PDCD1* and IFNg transcription and production in CD4^+^CD27^+^CD45RO^+^ memory T cells. Immune modulation impacted metabolic activity and synergized with the effect of histone acetylation on insulin responsiveness in RA patients.

**Conclusions:** RA synovia is enriched with the metabolic active *BIRC5*^hi^CD4^+^ T cell clusters. The metabolic activity of these cells is histone acetylation-dependent and mediates insulin effects through the H3K27ac-survivin epigenetic mechanism. Increasing plasma insulin levels when combined with insulin sensitivity, can be protective in RA dearmoring effector T cell function. Hence, increasing the insulin sensitivity by enabling histone acetylation presents a reasonable interventional goal to restore immune cell homeostasis in RA.

## Introduction

During chronic inflammation, hyperinsulinemia and insulin resistance is frequently found in rheumatoid arthritis (RA) patients (1), though the association with development of type 2 diabetes (T2D) is tenuous. Insulin resistance causes deep changes in epigenetic regulation of inflammatory genes such as *IL-6*, *TNF*, *IL-1β*, which promotes propagation of autoimmune disease including RA (2). Our studies in experimental RA demonstrated that induction of insulin resistance has dual effects. It alleviates inflammation and joint damage (3), while supporting autoimmunity and causing autoantibody production (4). This makes the understanding of insulin signalling central for RA pathogenesis. Investigation of RA patients (5,6) suggested that IR is a characteristic feature of a pathogenic subtype of T cells, amenable to targeting by anti-rheumatic drugs.

Insulin resistance implies the phenomenon in which cells are resistant to glucose uptake following insulin stimulation. Consequently, hyperglycaemia occurs, forcing further insulin production. In canonical T2D, excess glucose facilitates production of reactive oxygen species (7) and minimizes the immunomodulating effects of glucose, resulting in glucose toxicity and development of chronic insulin resistance. Activated cell function requires the tight regulation of several metabolic enzymes, such as pyruvate production via histone acetylation (8). Energy-demanding functions redirect cell metabolism into the pentose phosphate pathway (PPP) from aerobic glycolysis. Intermediates from pyruvate decarboxylation or PPP funnel into the TCA cycle, which generates acetyl Co-A and NADPH, activating the electron transport cycle for deriving energy through oxidative phosphorylation (OxPhos) (**Figure 1A**). Functional deficiency in any of the enzymes regulating these metabolic pathways changes the balanced flow of glucose utilization and sustains the pro-inflammatory or anti-inflammatory activity of a cell.

**Figure 1.**
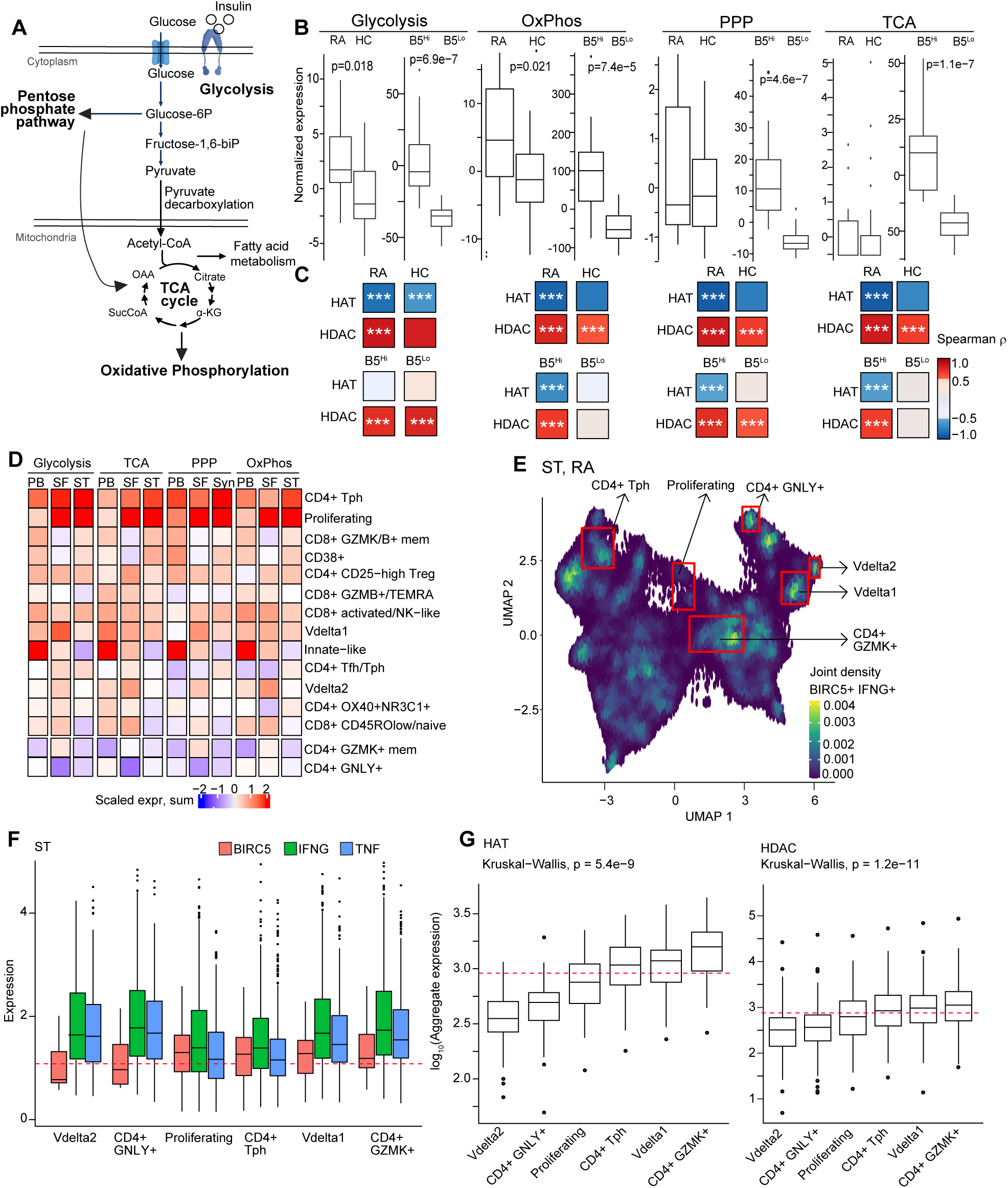
BIRC5^hi^CD4^+^ cells pinpoint migratory active CD4^+^ T cell clusters in blood and synovial tissue. A. Cartoon of glucose metabolism. The investigated metabolic pathways are underlined. B. Box plots of the normalized expression sum of genes acting within metabolic pathways in CD4^+^ cells of RA (n=11), healthy controls (HC, n=57), in RA CD4^+^ cells split by mean expression of *BIRC5* (B5) gene in BIRC5^hi^ and BIRC5^lo^ (n=24). Glycolysis (GO:0006096); pentose phosphate pathway (PPP, GO:0006098); oxidative phosphorylation (OxPhos, GO:0006119 and GO:0022900); tricarboxylic acid cycle (TCA, GO:0006099). Wilcoxon unpaired p-values are indicated. C. Heatmap of Spearman correlation of the HAT and HDAC enzyme genes and the corresponding metabolic pathways in CD4^+^ cells of RA and HC; and CD4^+^ cells split into B5^Hi^ and B5^Lo^. Asterisks indicate p-values. * < 0.05, ** < 0.01, *** <0.001 D. Heatmap of scaled total expression of the metabolic pathway genes in peripheral blood (PB), synovial fluid (SF), and synovial tissue (ST) of RA patients. Metabolically active T cell clusters are red and inactive clusters are blue. E. Uniform manifold approximation and projection (UMAP) map of joint *BIRC5* and *IFNG* density expression in T cell clusters in single-cell transcriptome of RA ST. *BIRC5^Hi^*CD4^+^ clusters are marked. F. Boxplot of gene expression of *BIRC5*, *IFNG*, and *TNF* in the *BIRC5^Hi^*CD4^+^ clusters in RA ST. G. Boxplot of HAT and HDAC enzymes gene expression in BIRC5^Hi^ clusters in RA synovial tissue. The red dotted line indicates the median expression of the non-BIRC5^Hi^ clusters. Kruskal-Wallis and pairwise Dunn test p-values adjusted with Bonferroni correction are indicated. * < 0.05, ** < 0.01, *** <0.001, **** <0.0001

Clinically, sensitivity to insulin is evaluated through blood and imaging tests. While these tests answer the question of how much insulin or glucose in the blood constitute abnormality, molecular indices to threshold insulin resistance and pertinent phenotypical characteristics defining a cell’s insulin sensitivity are still lacking.

Insulin has been shown to affect chromatin accessibility acting through modulation of histone modifications, activation of chromatin remodelling complexes, and recruitment of transcription factors and co-activators (9). The importance of histone acetylation for insulin sensitivity is stressed by the fact that histone deacetylase (HDAC) inhibition has been investigated for treatment for T1D and T2D (10) and in experimental arthritis (11). In mice, it has been observed that glucose restriction contributes to decrease in T cell effector function by altering histone methylation enzyme EZH2 (12). Further, HDAC5 activity is known to suppress GLUT4 translocation, thus affecting glucose uptake, while inhibition of HDAC3 activates PPARγ promoter and therefore development of insulin resistance (13). HDAC inhibitors are effective in releasing insulin resistance in mice while clinical trials in T2D showed mixed results (14,15). In RA synovium, opposing findings have been documented (16,17), where both hyperacetylation through reduced HDAC activity and reduced acetylation through increased HDAC activity were empirically attributed to RA pathogenesis. Presently, it remains unclear how the histone acetylation levels contribute to the development of pathogenic T cells in RA.In human and mouse CD4^+^ cells, activation-induced glycolysis depends on histone acetylation (8). Upon activation, CD4^+^ T cells utilize intermediates from the glycolytic pathway to supplement their energy needs regardless of oxygen levels. Survivin, an oncogenic protein central to RA pathogenesis, coordinates metabolic adaptation of T cells by interacting with chromatin modifying complexes rendering insulin resistance and DNA damage response (18–20). An association between cytotoxic elimination of survivin expressing pathogenic cells with high levels of survivin in serum has been demonstrated (21). High serum levels of survivin are known to predict development of RA (22,23) and are sensitive indicators of drug-resistant disease quickly progressing to joint damage (24,25). Negative seroconversion of survivin is associated with a robust control of the disease activity (24,26). Recently, we proposed a novel survivin-dependent mechanism which coordinated glycolytic switch of CD4^+^T cells in favor of pentose phosphate pathway to maintain activity of IFNγ-producing CD4^+^T cells (5). Insulin has a strong immunomodulating effect by suppressing IFNγ production by CD4^+^T cells. The initial anti-inflammatory properties of insulin signaling render senescence and elimination of the glucose-consuming effector T cells, which could be overcome under the conditions of insulin resistance, thereby promoting accumulation of the activated effector T cells.

In this study, we investigated how co-deposition of H3K27ac and survivin on regulatory chromatin contributes to insulin response in CD4^+^ cells. We found that co-deposition of survivin-H3K27ac guides insulin responsiveness and defines metabolically active CD4^+^ T cell clusters in blood and synovial tissue of RA patients. Insulin stimulation remarkably increased H3K27ac deposition, implying HDAC inhibition, thereby promoting a memory phenotype and suppressing T cell exhaustion. Transcriptional response to insulin elicited by survivin-H3K27ac deposition is only partly targetable by modern anti-rheumatic drugs.

## Methods

### Human material

This study used biological material collected from patients with rheumatoid arthritis (RA) and healthy controls summarized in **Supplementary Figure 1A**. We used three independent patient cohorts, which were collected at the Rheumatology Clinic of Sahlgrenska University Hospital, Gothenburg, Sweden. Cohort one was collected between 10 January 2022, and 14 March 2023. It consisted of 11 non-diabetic treatment naive RA patients and 57 non-diabetic subjects having no rheumatic diseases (healthy controls, HC), randomly selected among the 1st-visit patients with musculoskeletal complains. Cohort two was randomly collected during November 2018 and consisted of 24 RA patients with established disease. Cohort three was collected between 7 October 2019 and 26 October 2020. It consisted of 56 RA patients with established disease. All RA patients fulfilled the 2010 EULAR/ACR classification criteria (27). Among the RA patients within cohort 3, 24 were treated with the Janus kinase inhibitors (JAKi). Twelve patients combined JAKi with methotrexate, four with other antirheumatic drugs, and one each with abatacept, tocilizumab, or sarilumab. Among the non-JAKi-treated patients, fourteen were treated with methotrexate monotherapy, ten with TNF inhibitors, and one with tocilizumab. Two of the JAKi-treated and three of the non-JAKi-treated patients used oral corticosteroids. 8 patients had neither methotrexate, JAKi, nor oral corticosteroids. None of the control subjects were using immune suppressive drugs or oral corticosteroids. All study persons gave their written informed consent to physical examination and blood sampling. This study was approved by the Swedish Ethical Review Authority and was registered at clinicaltrials.gov with ID NCT03449589, NCT03444623.

For the *ex vivo* experiments and insulin treated RNAseq, cells from additional healthy non-diabetic subjects (5 female, 16 males; age 43.5 ± 14.5 years) were used.

### Isolation and stimulation of cells

Human peripheral blood mononuclear cells (PBMC) were isolated from the venous heparinized peripheral blood by density gradient separation on Lymphoprep (Axis-Shield PoC As, Dundee, Scotland). PBMC cultures (1 x 10^6^ cells/mL) were stimulated with ConA and LPS for 48h. After 24 hours the cultures were supplemented with insulin 25nM and/or the pan-HDAC inhibitor (HDACi) valproate, 50 µg/mL (stock 200 mg/mL, Ergenyl, Sanofi, Paris, France) as indicated. Supernatants were collected for cytokine measure and cells were analyzed by flow cytometry.

CD4^+^ cell cultures (1.25×10^6^ cells/mL) were prepared from fresh PBMC cultures by positive selection (Invitrogen, 11331D) in RPMI medium (Gibco, Waltham, Massachusetts, USA) containing 50μM β-mercaptoethanol (Gibco), Glutamax 2 mM (Gibco), gentamicin 50 μg/mL (Sanofi-Aventis, Paris, France) and 5% fetal bovine serum (Sigma-Aldrich) at 37°C in a humidified 5% CO_2_ atmosphere. For ChIPseq analysis, CD4^+^ cells were stimulated with concanavalin A (ConA, 0.625 μg/mL, MP Biomedicals), and lipopolysaccharide (LPS, 5 μg/mL, Sigma-Aldrich). For RNAseq and qPCR, cells were cultured in wells coated with anti-CD3 antibody (0.5 μg/mL; OKT3, Sigma-Aldrich, St. Louis, MO, USA), and treated with insulin (0 and 10 nM, Humalog 100 U/mL, Eli Lilly, Indianapolis, IN, USA) for 72h.

### Flow cytometry

At the end of incubation, cells were collected. The adherent cells were detached using 1 mM ice cold EDTA for 15 minutes. The adherent and non-adherent cells were pooled, and cells were suspended in phosphate buffered saline containing 2% fetal calf serum, 0.1% sodium azide and 1 mM EDTA. Non-specific binding was blocked with human γ-globulin (Beriglobin, CSL Behring, Melbourne, Australia), thereafter monoclonal mouse anti-human antibodies CD4-Brilliant Violet 510 (Clone SK3, Beckton Dickinson, Franklin Lakes, NJ, USA), CD14-PE (Clone M5E2, BD), CD45RO-APC (clone UCHL1, BD), CD27-FITC (clone L128, BD) and PD1-Brilliant Violet 421 (clone EH12.1, BD) were added. All antibodies were diluted in FACS buffer (Beckton Dickinson) to optimal concentrations. The stained cells were collected 100 000 events/sample in the clinical analyzer FACS-Lyric (Beckton Dickinson) and analyzed with Flow-Jo software version 10.10 (Beckton Dickinson). Cell populations were gated using fluorochrome minus one staining controls.

### Cytokine measurements

Cytokine levels were measured by sandwich ELISAs for IFNγ (DY285B), IL-4 (DY204), and IL13 (DY213) (R&D Systems, Minneapolis, MN, USA), according to the manufacturer’s instructions.

### Transcriptional sequencing (RNA-seq)

RNA of CD4^+^ cells was prepared using the Norgen Total RNA kit (17200 Norgen Biotek, Ontario, Canada). Quality control was done by Bioanalyzer RNA6000 Pico on Agilent2100 (Agilent, Santa Clara, CA, USA). Deep sequencing was done by RNA-seq (Hiseq2000, Illumina) at the core facility for Bioinformatics and Expression Analysis (Karolinska Institute, Huddinge, Sweden). Raw sequence data were obtained in Bcl-files and converted into fastq text format using the bcl2fastq program from Illumina.

### Chromatin immunoprecipitation and sequencing (ChIP-seq)

For ChIP-seq analysis, three CD4^+^ cell cultures were stimulated with ConA and LPS for 24 h. The cells were cross-linked and lysed with the MAGnify Chromatin Immunoprecipitation System (492024, Applied Biosystems, ThermoFisher Scientific). After sonication, cellular debris was removed, and DNA material was pooled. After preclearing, 1% of the sample was saved as an input fraction and used as background binding. Pre-cleared chromatin was incubated with 2 μg of anti-survivin (ab192675, Abcam, Cambridge, UK), or anti-H3K27ac (C15410196, Diagenode). The immune complexes were washed, the cross-links were reversed, and the DNA was purified as recommended. The quality of purified DNA was assessed with TapeStation (Agilent, Santa Clara, CA, USA). DNA libraries were prepared with ThruPLEX (Rubicon) and sequenced with a Hiseq2000 sequencing system (Illumina). Bcl-files were converted and demultiplexed to fastq with bcl2fastq (Illumina).

### Immunohistochemistry

Human monocytic cell line THP1 (TIB-202, ATCC, Manassas, VA, USA) were seeded 10^6^/mL on glass chamber slides (ThermoScientific) precoated with poly-L-lysine (Sigma-Aldrich, Saint Louis, MO, USA). Cells were treated with insulin 0nM, 10nM and 25nM (Humalog 100 U/mL, Eli Lilly) for 24h. At harvest, cells were fixed with 4% buffered formalin for 10 minutes, thereafter blocked and permeabilized for 3 hours with 3% normal goat serum and 1% Triton-X100. Primary antibodies against H3K27ac and isotype controls were diluted in blocking buffer and the slides were incubated over night at 4°C. This was followed by Alexa-fluor conjugated secondary antibody donkey-anti-rabbit AF647 (Invitrogen A31573) for 2 hours at room temperature. Autofluorescence was blocked with 0.5% Sudan Black B (Sigma-Aldrich) in 70% ethanol for 20 minutes at room temperature. Nuclei were stained with Hoechst 34580 (NucBlue Live Cell Stain; Thermo Fisher Scientific, Waltham, MA, USA) for 20 minutes and mounted with ProLong Gold antifading mounting reagent (Invitrogen).

### Confocal imaging and analysis

Fluorescence microscopy was performed using the confocal imaging system Leica SP8 (Leica Microsystems, Wetzlar, Germany) with sequential acquisition using a 40x oil objective and up to 10x digital zoom. The images were acquired at high resolution (1.5x digital zoom) viewing 20-40 nuclei per image. Within each sample, H3K27ac-positive foci were enumerated in 2-3 images resulting in 44-63 nuclei per treatment. Images were analyzed with ImageJ version 2.9 within 8-bit composite images (28). Threshold was adjusted to optimize identification of positive spots. Nuclear area was defined by thresholding the Hoechst (blue) image and exporting the results to ROI.

## Bioinformatics analysis

### RNA-seq analysis

Mapping of transcripts was done using Genome UCSC annotation set for hg38 human genome assembly. Analysis was performed using the core Bioconductor packages in R-studio v. 4.4.1. Differentially expressed genes (DEG) between the samples were identified using DESeq2 (v.1.44.0) with Benjamini-Hochberg adjustment for multiple testing.

### ChIP-seq analysis

The fastq sequencing files were mapped to the human reference genome (hg38) using the STAR aligner (29) with default parameters apart from setting the alignIntronMax flag to 1 for end-to-end mapping. Quality control of the sequenced material was performed by FastQC tool using MultiQC v.0.9dev0 (Babraham Institute, Cambridge, U.K.). Peak calling was performed using the HOMER (30) findPeaks command, with 1 tag per base pair counted (-tbp 1). For peak calling in histone ChIP-seq, the option -style histone was used to find broad regions of enrichment. Peaks were filtered for the histone H3 antibody or survivin antibody IP fraction and unprocessed DNA (Input), which is a generally accepted normalization approach to identify protein-specific enrichment of DNA interaction areas (31). A set of peaks with enrichment versus surrounding region and Input (adjusted p < 10e^−5^) was identified and quantified separately for each sample. Peaks were annotated with HOMER software in standard mode to the closest TSS. Peaks with overlapping localization by at least one nucleotide were merged and further on referred to as one peak. To quantify strength of binding and maintain consistency of comparison in the histone H3 samples and survivin sample, peak score was calculated by the position adjusted reads from initial peak region.

### Identification of peak colocalization

The R package ChIPpeakAnno (32), version 3.38.1 was used to identify colocalization of the control ChIP-seq peaks of survivin and the histone H3K27ac peaks. The function ‘findOverlapsofPeaks’ was used, with parameters restricting the maximum gap between peak ranges to zero, indicating a minimum of one bp overlap, and connected peak ranges within multiple groups as ‘merged’. The resulting set of chromatin regions represented the genome locations possessing concomitant deposition of survivin and H3K27ac.

### Genomic regulatory element colocalization and overlap with TF target genes

Experimentally confirmed candidate regulatory elements (RE) was obtained through the GeneHancer database version 5.9 by request (33). The BED files of RE were combined with BED files of gene bodies and 2kb upstream promoters of hg38 to obtain an integrated list of genomic regulatory elements. Using the ChIPpeakAnno parameters mentioned above, chromatin regions possessing colocalized survivin and H3K27ac were overlapped with the genomic regulatory elements. Further, the entire list of the genes connected to the genomic regulatory elements was retrieved used for further downstream analysis.

### Genes relating to metabolic pathways

For analysis of metabolic activity, we used the Gene Ontology database (**34**) to retrieve the genes annotated to the four metabolic pathways: Glycolysis, GO:0006096; Pentose Phosphate Pathway (PPP), GO:0006098; Tricarboxylic Acid (TCA) cycle, GO:0006099; and Oxidative phosphorylation (OxPhos), GO:0022900 and GO:0006119. To analyze histone acetylation activity, we retrieved the transcription levels of the genes annotated to histone acetyl transferase enzymes (HATs), *ATF2, BRCA2, BRD1, BRPF1, BRPF3, CLOCK, CREBBP, EP300, GTF3C4, HAT1, ING3, ING4, JADE1, JADE2, KAT2B, KAT5, KAT6A, KAT6B, KAT7, KAT8, MSL3, NAA50, NAA60, NCOA1, NCOA3, PHF10, TADA2A, TAF1, TAF10, TAF9, USP22*; and genes related to histone deacetylases (HDACs), GO:0000118.

### Single cell transcriptome (scRNAseq) analysis

To validate the insulin responsiveness gene signature obtained through bulk transcriptomics, we used the single cell transcriptome datasets of synovial tissue (**35**), synovial fluid and PBMC of RA patients (n=8) (**36**), T2D (n=7) and healthy controls (n=3) (**37**). Using the 24 T cell clusters of RA synovia (94,046 cells) and the logNormalized matrix of gene counts, we performed a pseudobulk aggregated expression of the metabolic pathways (see above) and aberrant insulin-responsive genes in each cluster. Using the cluster annotation of RA synovia, we mapped (using *FindTransferAnchors* and the *MapQuery* functions in Seurat) and annotated cell labels in the RA PBMC single cell dataset after merging of the PBMC samples and similar pre-processing steps as for the RA synovia. Similarly, single cell transcriptome of PBMC from type-2 diabetes and healthy controls were retrieved (**37**), and T cell clusters were mapped. To assess enrichment, T cell clusters that showed fewer than 3% of the total cell population in any of the single cell datasets were removed from the analysis.

### Data analysis and visualization

Statistical analysis was performed using R-studio (version 4.4.1). Heatmaps were visualized using the R package ComplexHeatmap (38) version 2.20.0. Analysis for the enriched biological pathways was done through the Gene Set Enrichment Analysis (GSEA) online tool from Broad Institute, UC San Diego (39,40).

### Data availability

ChIPseq datasets for CD4^+^ samples immunoprecipitated with survivin and H3K27ac is available in NCBI GEO with accession GSE282301. Transcriptome sequencing data of insulin-stimulated CD4^+^ cells is deposited in NCBI GEO with accession GSE282515. Transcriptome sequencing data of CD4^+^ cells from RA patients and healthy controls is deposited in NCBI GEO with accession GSE282517. Transcriptome sequencing data of HDACi-treated CD4^+^ cells is deposited in NCBI GEO with accession GSE132053. Single-cell sequencing data for CD4^+^ cells from blood and synovial fluid (36), RA synovial tissue (35) are publicly available. PBMCs isolated from T2D is accessible in NCBI GEO with accession GSE165816 (37). For assessment of the effect of anti-rheumatic drugs on transcriptome of CD4+ cells we used paired transcriptome data of CD4^+^ cells isolated from untreated RA patients (n=5) and treated with JAKi (n=27), CTLA4 fusion protein abatacept (n=22 (41) and n=14 (42)), folic acid antagonist methotrexate (n=28 (43)) and IL6 receptor inhibitor tocilizumab (n=6 (44)). The biological material used in this study is summarized in **Supplementary Figure 1A**.

## Results

### Glucose utilization transcriptome is dysregulated in BIRC5^hi^ CD4^+^ cells in rheumatoid arthritis

We investigated the metabolic profile of CD4^+^ cells by studying the expression of genes involved in the canonical metabolic pathways (**Figure 1A**) in patients with rheumatoid arthritis (RA, n=11) and healthy controls (HC, n=57). This analysis revealed that CD4^+^ cells in RA had significantly activated glycolysis, and oxidative phosphorylation (OxPhos) pathways, while pentose phosphate pathway (PPP), and tricarboxylic acid (TCA) cycle were similar to HC (**Figure 1B**). In line with survivin’s role in mediating proinflammatory effects in CD4^+^ cells (5), we found that high expression of the *BIRC5* gene coding for survivin was associated with activation of all metabolic pathways in the CD4^+^ cells (**Figure 1B**). Acetylation of lysine at position 27 of histone H3 tail (H3K27ac) is controlled by the histone acetyltransferase (HAT) enzymes, which deposit acetylation of histone H3 tail to initiate gene transcription, while histone deacetylases (HDAC) abolish gene transcription by removing histone acetylation. Glucose metabolism via insulin signaling in both T2D and in RA is regulated by histone acetylation (9,10). Therefore, we investigated the correlation of HAT and HDAC enzymes to the metabolic profile of CD4^+^ cells between RA and HC. HDAC enzymes were positively correlated to the metabolic profile in RA conditions, while HAT enzymes were inversely related (**Figure 1C**). The correlation between the metabolic genes and HAT/HDAC enzymes was more pronounced in CD4^+^ cells with high *BIRC5* expression, where positive association to HDAC enzymes was seen in all metabolic pathways (**Figure 1C**).

Next, we aimed to identify the metabolically active CD4^+^ cell subsets present in blood and the joints of RA patients to characterize the clusters that contribute to chronic inflammation. Mapping specific markers of the 24 T cell clusters (45), we investigated the single-cell transcriptome of CD4^+^ cells in blood and synovial fluid of 8 RA patients (36) and extended it further into synovial tissue of 82 RA patients with active disease (45).

In RA patients, this approach identified a subset of the metabolic active T cell clusters, characterized by high transcription of enzymes mediating the metabolic pathways of glycolysis, PPP, TCA, and OxPhos (**Figure 1D**). Among these metabolic active T cell clusters, co-expression of *BIRC5* and *IFNG* in RA synovia pinpointed the clusters of the PD1^+^CXCL13^+^ Tph cells, PTTG1^+^TUBA1B^+^ proliferating cells, CD4^+^GNLY^+^GZMB^+^CD45ROneg cells, and CD4^+^GZMK^+^ memory cells, as well as Vdelta1 and Vdelta2 cells (**Figure 1E**). Notably, aggressive nature of these cell clusters is revealed by a strong correlation between expression of the metabolic pathway genes and the markers defining these clusters (**Supplementary Table T1**), and, also, the strong upregulation of expression of the cluster markers in blood BIRC5^hi^CD4^+^ cells of RA patients (**Figure 1E**). Interestingly, the metabolic active clusters of the peripheral helper T cells (Tph), proliferating, CD4^+^GZMK^+^, CD4^+^GNLY^+^ and Vdelta cells also accumulated in the RA synovial tissue and had high expression of *BIRC5, IFNG* and *TNF* genes (**Figure 1D, F**), suggesting a migratory destiny of these cell populations to RA joints. In these *BIRC5*^hi^ *IFNG*^hi^ T cell clusters, we also observed variable expression of HAT and HDAC enzymes. Both HATs and HDACs were high in the synovial CD4^+^GZMK^+^ cluster, while the CD4^+^GNLY^+^ and Vdelta2 cell clusters had the HAT and HDAC expression significantly below the median expression of the non-BIRC5^Hi^ clusters (**Figure 1G**). The other T cell populations were metabolically dormant, both in synovial tissue and in blood, suggesting their accessory role in chronic inflammation in RA. Inhibition of HDAC in CD4^+^ cells *ex vivo* significantly suppressed the markers of these clusters (**Supplementary Figure S1B**), while insulin had a prominent suppressive effect on the markers of the CD4^+^Tph cluster. Increasing plasma insulin levels correlated to the scaled normalized expression of the insulin signaling genes *INSR, IRS1, IRS2* (**Supplementary Figure S1C**).

Together, our analysis revealed that the increased expression of metabolic pathway genes was closely linked to the high expression of *BIRC5* and histone acetylation enzymes in CD4^+^ cells in RA. Co-expression of *BIRC5* and *IFNG* demarcated the metabolically active CD4^+^ clusters migrating into RA tissues.

### Insulin-dependent transcription in *BIRC5*^hi^CD4^+^ cells promote chromatin remodeling and immune response

To understand how *BIRC5^hi^*CD4^+^ cells respond to insulin, we first performed sequencing of chromatin of CD4^+^ cells immunoprecipitated using survivin and H3K27ac antibodies. Survivin-bound chromatin pinpointed H3K27ac deposition in 2452 genomic regions (**Figure 2A**). Notably, 68% (n=1668) of these genomic regions localized within *cis*-regulatory elements (*cis*-RE) annotated as enhancers (29.3%), promoters (24.1%), and gene bodies (46.6%). Among the genes connected to *cis*-RE containing deposition of survivin and H3K27ac, we identified a substantial part of those involved in the metabolic pathways (**Figure 2B**). To study effect of insulin, HDACi and survivin on the metabolic pathway genes connected to *cis*-RE containing survivin and H3K27ac deposition, we performed DESeq2 statistics on transcriptome data of primary healthy CD4^+^ cells treated with insulin (n=6), HDACi romidepsin (n=3 (46)) and survivin inhibitor YM155 (n=4 (5)). HDACi differentially affected the expression of about 40% of the metabolic pathway genes overlapping with the H3K27ac-survivin controlled in about 10% and resulted in activation of glycolysis and PPP (**Figure 2C**). Neither insulin stimulation nor survivin inhibition had an immediate effect on transcription of the metabolic pathway genes in CD4^+^ cells. In line with the earlier observation of positive correlation between the expression of metabolic genes and HDAC in *BIRC5*^hi^CD4^+^ cells (**Figure 1C**), *BIRC5* was significantly repressed by HDACi (**Figure 2D**), and survivin inhibition raised the levels of HAT and HDAC where the change difference was significant for HATs (**Figure 2E**). Surprisingly, CD4^+^ cells treated with HDACi showed increased transcription of HDAC enzymes and suppression of HAT enzymes (**Supplementary Figure S1D**), while insulin did not have a significant effect.To investigate the insulin-dependent transcription enacted by survivin-H3K27ac chromatin deposition, we first identified the genes connected to survivin-H3K27ac deposition that showed upregulation in *BIRC5*^hi^CD4^+^ cell clusters migrated to RA synovia. Further, we subset this list to the HAT/HDAC-correlated and insulin-dependent genes obtained by continuous regression using plasma insulin levels in the RA-HC cohort (**Figure 2F**). These genes represent the connection between histone acetylation and insulin-dependent transcription in *BIRC5^hi^*CD4^+^ cells. We found that insulin stimulated CD4^+^ cells showed major upregulation of the H3K27ac-survivin deposition controlled genes including immune-related processes of lymphocyte activation (GO:0046649: FDR=2.21e^-13^; 34 genes), cell surface receptor signaling (GO:0002768: FDR=2.50e^-10^; 21 genes), and cytokine signaling (R-HSA-1280215: FDR=2.80e^-6^; 23 genes) (**Figure 2G, Supplementary Table T2**). Pathways suppressed by insulin included the H3K27ac-survivin deposition controlled genes with chromatin-intrinsic functions acting cell cycle (GO:0007049: FDR=6.8e^-31^; 88 genes), DNA damage response (GO:0006974: FDR=1.24e^-22^; 56 genes), chromosome organization (GO:0051276: FDR=1.28e^-17^; 42 genes), and apoptosis (GO:0006915: FDR=4.99e^-15^; 69 genes).

**Figure 2.**
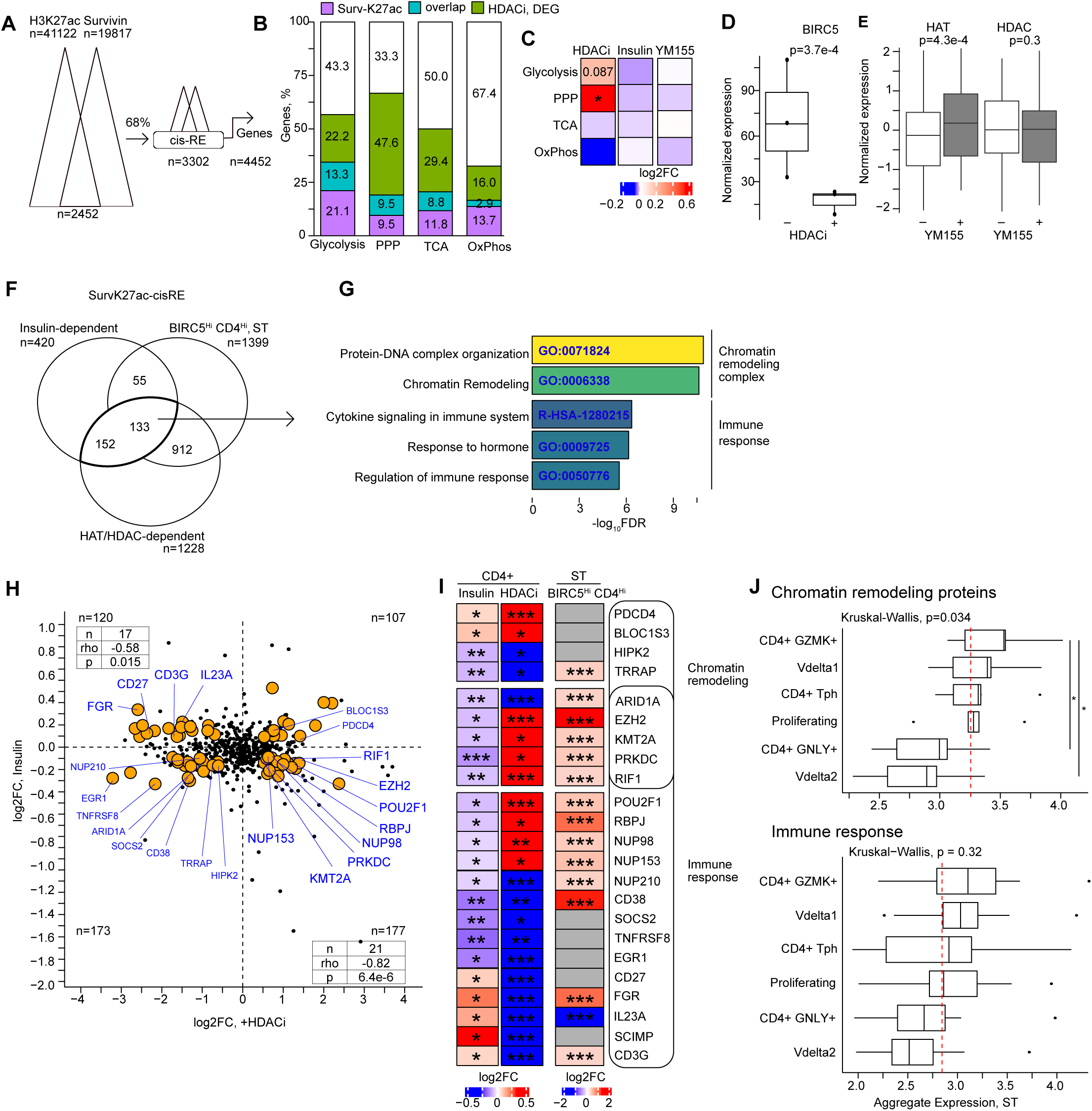
Insulin stimulation suppressed chromatin remodeling and promoted immune response regulated by survivin and H3K27ac. A. Cartoon of analysis strategy for survivin and H3K27ac deposition in cis-regulatory elements (*cis*-RE) of human genome. B. Stacked bar plot of percentage of genes connected to *cis*-RE containing survivin-H3K27ac (purple), differentially regulated by HDAC-inhibitor romidepsin (green), and the overlap (blue). Glycolysis (GO:0006096); pentose phosphate pathway (PPP, GO:0006098); oxidative phosphorylation (OxPhos, GO:0006119 and GO:0022900); tricarboxylic acid cycle (TCA, GO:0006099). Percentage numbers are mentioned inside the bars. C. Heatmap of gene expression change of metabolic pathway enzymes, by log2FC, in CD4^+^ cells treated with HDAC inhibitor romidepsin, insulin, and with survivin inhibitor YM155. Asterisks indicate paired Wilcoxon test p-values * < 0.05, ** < 0.01, *** <0.001 D. Boxplot of normalized expression of *BIRC5* gene in CD4^+^ cells treated with HDAC inhibitor romidepsin. Paired Wilcoxon test p-values are indicated. E. Boxplot of summarized expression of HAT and HDAC genes in CD4^+^ cells after YM155 treatment. Wilcoxon p-values are shown. F. Venn diagram of insulin-dependent genes correlated to histone acetylation enzymes HAT/HDAC and genes connected to cis-RE containing survivin and H3K27ac deposition. G. Bar plot of the enriched pathways of the intersection genes in Figure 2F. H. Scatterplot of expression change (by log2FC) after insulin and HDAC inhibitor treatment of CD4^+^ T cells. Genes differentially expressed after treatment with insulin and HDAC inhibitor are highlighted yellow. Spearman correlation of gene expression change after insulin stimulation and HDAC inhibition is shown in the table. Number of genes in each quadrant is indicated. I. Heatmap of gene expression change in enriched pathways (left) and T cell markers (right), by RNA-seq log2FC, in CD4^+^ T cells stimulated with insulin in vitro (n=6), after HDAC inhibition (n=3), and in BIRC5^Hi^CD4^+^ clusters in RA synovia. Asterisks indicate Wilcoxon paired test p-values, *< 0.05, **< 0.01, ***< 0.001 J. Boxplot showing aggregate expression of genes involved in chromatin remodeling and immune response in BIRC5^Hi^ CD4^+^ T cell clusters in synovial tissue of RA patients. The red dotted line indicates the median expression of the non-BIRC5^Hi^ clusters. Kruskal-Wallis and pairwise Dunn test p-values adjusted with Bonferroni correction are indicated. * < 0.05, ** < 0.01, *** <0.001, **** <0.0001

### Survivin-H3K27ac deposition synergize insulin-dependent effect to promote the memory T cell clusters in RA

To further define insulin’s effect on acetylation- and survivin-dependent transcription, we subjected CD4^+^ cells to insulin stimulation *ex vivo* and profiled their transcriptome through RNAseq. Stimulation with insulin altered the transcription of 796 DEG (**Supplementary Table T3**), 195 of which (24.5%) were connected to the *cis*-RE bound H3K27ac-survivin. By filtering the DEG sensitive to insulin stimulation *ex-vivo* and the DEG sensitive to increasing plasma insulin concentration in the RA-HC cohort, on the genes connected to H3K27ac-survivin *cis*-RE, we obtained the list of 585 insulin-responsive genes (**Supplementary Figure S1E, Supplementary Table T4**).

Among those insulin-responsive genes, we found a strong inverse correlation between insulin stimulation and HDAC inhibition (**Figure 1H**) i.e., genes activated by HDACi were suppressed by insulin (rho=-0.82, p=6.4e-6), and *vice versa*, the genes suppressed by HDACi were activated by insulin (rho=-0.58, p=0.015). Together, this suggested that effect of insulin was mediated by histone deacetylation. Prominent among the genes transcriptionally controlled by insulin and HDACi were *CD27, CCR7, SELL, S1PR1, IL23A, RBPJ, POU2F1, CD3G, ZNF683/HOBIT* representing a memory phenotype of CD4^+^ T cells, and *EZH2, ARID1A, KMT2A,* and *PRKDC* genes important for reorganizing the histone H3 methylation landscape of chromatin (**Figure 2I**). Additionally, insulin and HDACi presented joint effect by suppressing the exhaustion check-point receptors *PDCD1, CTLA4, CD38,* and *TIGIT* (**Figure 2I**).

Analyzing the insulin-responsive genes in the metabolically active BIRC5^hi^CD4^+^ cell clusters in RA synovial tissue, we found upregulated expression of both chromatin remodeling and immune response genes connected through *cis*-RE to H3K27ac-survivin co-deposition (**Figure 2I**). The memory Tph and GZMK^+^ clusters had high expression of the exhaustion check-point receptors *PDCD1*, *CTLA4*, *TIGIT*, and low/no expression of the memory receptors *CD27, CCR7, S1PR1, and SELL* (**Figure 3B, 3C**). The memory phenotype of these clusters is supported by high protein level of the memory CD45RO receptor in relation to the protein level of CD45RA, while the CD45RO/CD45RA ratio was low in the Vdelta1 and GNLY^+^ clusters (**Figure 3A**). Aggregate expression of the chromatin remodeling proteins *EZH2*, *KMT2A*, *RIF1*, *PRKDC*, *ARID1A* and *TRRAP*, was significantly different between the BIRC5^hi^CD4^+^ clusters, where the memory CD4^+^GZMK^+^ and CD4^+^Tph cell cluster with high expression of HAT/HDAC enzymes had also significantly higher expression of chromatin remodeling genes compared to the CD4^+^GNLY^+^ and Vdelta2 clusters where expression of HAT/HDAC was low (**Figure 2J, Figure 1G**). These observations make the metabolic active BIRC5^hi^CD4^+^ clusters an obvious target for insulin and HDACi targeted interventions.

**Figure 3.**
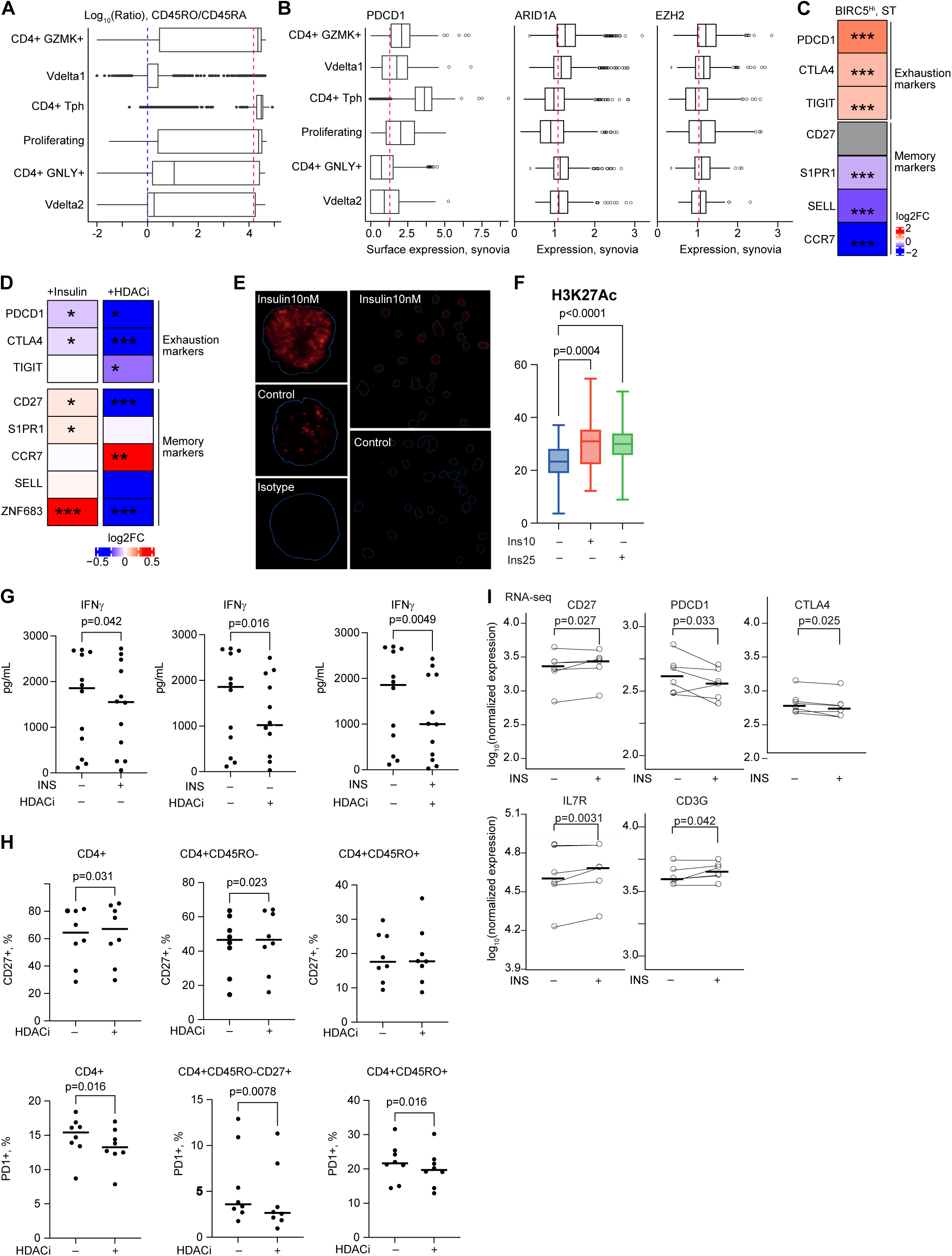
Insulin stimulation and HDAC inhibition suppress exhaustion and promote memory phenotype in CD4^+^ T cells. A. Boxplot of CD45RO/CD45RA ratio in BIRC5^Hi^CD4^+^ cell clusters in RA synovial tissue. B. Boxplot of normalized expression of T cell markers in BIRC5^Hi^CD4^+^ clusters in RA synovial tissue. C. Heatmap of gene transcription difference between BIRC5^Hi^CD4^+^ clusters and the remaining T cell clusters in RA synovial tissue, by log2 fold change (FC). D. Heatmap of gene transcription difference in CD4^+^ cells stimulated with insulin and HDAC inhibitor romidepsin, by log2FC. Asterisks indicate Wilcoxon paired test p-values, *< 0.05, **< 0.01, ***< 0.001 E. Confocal microscopy images of H3K27ac (stained red) in nucleus of THP1 cells stimulated with control or insulin. F. Box plots of H3K27ac foci quantification in THP1 cells before and after stimulation with insulin. Wilcoxon test p-values are shown G. Scatterplot of IFNγ levels in supernatants of CD4^+^ cells stimulated with insulin and/or HDAC inhibitor valproate. Solid line indicates median. Wilcoxon test p-values are shown H. Scatterplot of percentage of CD27 and PD1 in CD4^+^ cells before and after treatment with HDAC inhibitor valproate, analyzed by flow cytometry. Wilcoxon test p-values are shown I. Scatterplot of RNA-seq normalized expression of memory T cell markers in control and insulin-stimulated CD4^+^ cells. Solid line indicates median. DESeq2 p-values are shown.

### Insulin stimulation and HDAC inhibition suppress T cell exhaustion in memory CD4^+^ T cells

To investigate the role of insulin and HDAC in development of the memory T cell phenotype, we first treated THP1 cell cultures with insulin or HDACi valproate. Exposure to insulin led to the accumulation of significantly higher numbers of H3K27ac^+^ loci in cell nuclei (**Figure 3E, 3F**). Analogously, treatment of THP1 cells with HDACi significantly increased deposition of H3K27ac in cell nuclei (*not shown*).

*Ex vivo* stimulation of CD4^+^ cells with insulin suppressed transcription of exhaustion receptors *PDCD1*, *CTLA4*, *TIGIT*, and activated the memory markers *CD27, CD3G, IL23A* and *S1PR1* (**Figure 3I**). In the next step, we performed flow cytometry in blood leukocytes treated with HDACi gating on CD4^+^CD14^-^ cells. We found that treatment with HDACi significantly reduced the surface expression of PD1 receptor on CD4^+^ cells by decreasing PD1^+^ subset within both the naïve CD4^+^CD45RO^-^ and the memory CD4^+^CD45RO^+^ cells, which justified a release of exhausted T cell phenotype (**Figure 3H**). Concordantly, HDACi increased expression of CD27 on the total CD4^+^ population and specifically on the CD4^+^CD45RO^-^ cells (**Figure 3H**). The reduction of the exhausted CD4^+^ cell subset was associated by lower production of protein IFNγ measured in the supernatants of insulin and HDACi treated cell cultures (**Figure 3G**).

These experimental findings evidence a significant impingement of insulin and HDACi on the memory and exhausted phenotype of CD4^+^ cells acting through control of histone acetylation-dependent gene transcription.

### Anti-diabetic treatment suppresses immune response in RA

To verify whether the identified insulin-responsive metabolic profile was associated with canonic insulin resistance of T2D, we mapped the 24 T cell clusters (45) into the single-cell transcriptome of blood mononuclear leukocytes of T2D patients (n=3) and healthy controls (HC, n=3) (37). This analysis demonstrated that the leukocytes of both T2D patients and HC contained only 11 of 24 T cell clusters present in RA patients (**Supplementary Figure S1F, Supplementary Table T5**). The proliferating T cell cluster of T2D patients was metabolic active.

To investigate if T2D and RA had similarities in the histone acetylation-dependent gene transcription, we performed pathway enrichment analysis on the differentially expressed genes in T2D vs HC and overlapped them with the pathways represented by the insulin-responsive genes correlated to HAT/HDAC. We identified 13 common pathways presented by 58 insulin-responsive genes (**Figure 4A, Supplementary Table T6**). Majorly, the enriched pathways were related to the immune response such as inflammatory response (GO:0006954: *RORA, FPR1, TNFRSF1A, IL10RA, LGALS1, ITGAL, GRN, LYZ*), and immune cell maturation such as lymphocyte activation (GO:0051249: *LGALS1, ITGAL, CD3E, ZFP36L1, RORA, CD3G, CD3D, CD8A, CD8B, DCAF12*), and leukocyte differentiation (GO:0002521: *LGALS1, CD3E, ZFP36L1, RORA, CD3G, CD3D, CD8A*) and, also, cytokine signaling in immune system (R-HSA-1280215: *RORA, FPR1, TNFRSF1A, IL10RA, NLRC5, PSMB3, DUSP6, TUBA4A, SLA*). Treatment of healthy CD4^+^ cells with HDACi significantly suppressed the expression of the genes common for T2D/RA T cells (**Figure 4B**).

**Figure 4.**
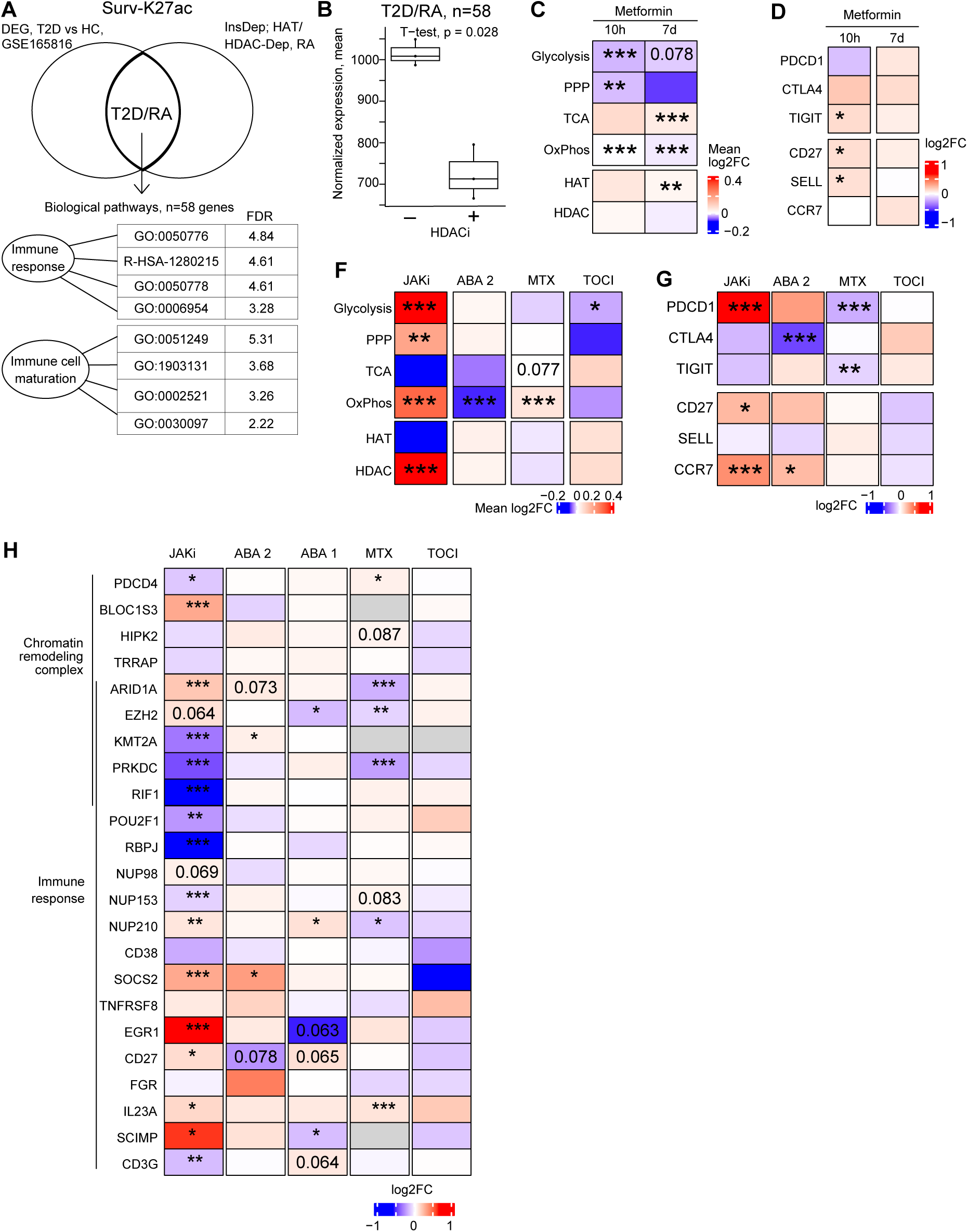
Modern anti-rheumatic drugs suppress pathogenic insulin-responsive gene signature in CD4^+^ cells of RA patients. A. Venn diagram of intersection of top enriched pathways of differentially expressed genes in T2D patients and insulin-dependent genes correlated to HAT/HDAC enzymes in CD4^+^ T cells of RA patients. Table shows the false discovery rate (FDR)-adjusted p-values of the enriched pathways B. Boxplot of normalized expression of insulin-dependent genes in the common T2D/RA pathways before and after treatment with HDAC inhibitor romidepsin. Unpaired T-test p-values are shown. C. Heatmap of expression difference of metabolic pathway genes in CD4^+^ cells of healthy controls treated with metformin short-term (10h), long-term (7 days), by RNA-seq log2 fold change (FC). Asterisks indicate Wilcoxon unpaired test p-values, *< 0.05, **< 0.01, ***< 0.001 D. Heatmap of T cell gene expression in CD4^+^ cells of healthy controls treated with metformin short-term (10h), long-term (7 days), by RNA-seq log2FC. Asterisks indicate Wilcoxon unpaired test p-values, *< 0.05, **< 0.01, ***< 0.001 E. Heatmap of expression sum for metabolic pathway enzymes gene in CD4^+^ cells of RA patients treated with JAKi (35 treated, 5 untreated), abatacept 1 (ABA 1, n=22, paired), methotrexate (MTX, n=28, paired), tocilizumab (TOCI, n=6, paired), by RNA-seq log2FC. Wilcoxon p-values are shown. *< 0.05, **< 0.01, ***< 0.001 F&G. Heatmap of normalized expression of T cell marker genes (F) and pathway-enriched insulin dependent genes (G) in CD4^+^ cells of RA patients treated with JAKi (35 treated, 5 untreated), abatacept 1 (ABA 1, Zenodo 8250013, n=22, paired), abatacept 2 (ABA 2, GSE121827, n=14, paired), methotrexate (MTX, n=28, paired), tocilizumab (TOCI, n=6, paired), by RNA-seq log2FC. Wilcoxon p-values are shown. *<0.05, **<0.01, ***<0.001 H. Heatmap of gene expression change in enriched pathways (left), by RNA-seq log2FC, in CD4^+^ T cells of RA patients treated with JAKi (35 treated, 5 untreated), abatacept 1 (ABA 1, Zenodo 8250013, n=22, paired), abatacept 2 (ABA 2, GSE121827, n=14, paired), methotrexate (MTX, n=28, paired), tocilizumab (TOCI, n=6, paired), by RNA-seq log2FC. Wilcoxon p-values are shown. *<0.05, **<0.01, ***<0.001

To study how improvement of insulin sensitivity affected the histone acetylation-dependent immune response, we analyzed the whole blood transcriptome of healthy individuals (n=25) who received short-term (10 hours) and long-term (7 days) metformin treatment (47). Expectedly, metformin suppressed the metabolic pathways in blood cells with both treatment regimens (**Figure 4C**). This suppression of the metabolic pathways was associated with downregulation of the H3K27 methyltransferase *EZH2* and *HIPK2,* but not other insulin-responsive chromatin remodeling genes. Among the immune response genes, metformin upregulated important mediators of T cell maturation *CD27, IL23A, RBPJ*, *CD3G, SELL and TIGIT* (**Supplementary Figure S1G**), thus promoting insulin-dependent effects and insulin sensitivity of CD4^+^ cells (**Figure 4D**). The observed improvement of T cell maturation in conjunction to metformin treatment could be a consequence of inflammation suppression in the metabolically active BIRC5^hi^CD4^+^ cells by downregulation of *CCL3, KLRD1, GNLY, GZMB, CCL4, TNF* genes (**Supplementary Figure S2A**), which represented the effector Vdelta1+2, CD4^+^GZMK^+^ and CD4^+^GNLY^+^clusters with low memory CD45RO/CD45RA ratio. In contrast, the HLA-DR genes representing the proliferating CD4^+^ cell cluster were activated by metformin treatment (**Supplementary Figure S2A**).

Taken together, our data revealed common pathways of the immune response in T2D and RA CD4^+^ cells that could be suppressed by metformin and HDACi treatment within BIRC5^hi^CD4^+^ clusters. Therefore, our results point towards the fact that interventions improving insulin sensitivity of CD4^+^ cells promote acquisition of T cell memory and mitigate their effector ability.

### Anti-rheumatic drugs alter metabolic profile of RA CD4^+^ T cells and address synovial BIRC5^hi^ pathogenic clusters

To assess if anti-rheumatic drugs influenced metabolic activity and insulin-responsive genes, we retrieved paired transcriptome data of CD4^+^ cells isolated from RA patients treated with JAKi (n=27 vs 5 untreated), CTLA4 fusion protein abatacept (n=22 (41) and n=14 (42)), folic acid antagonist methotrexate (n=28 (43)) and IL6 receptor inhibitor tocilizumab (n=6 (44)) (**Supplementary Table S1A**). We found that JAKi had significant activating effect on HDACs and the genes of glycolysis, and PPP pathways in CD4^+^ cells. Effects on the OxPhos pathway responsible for establishment of T cell memory (48,49), were common for methotrexate and JAKi, which transcriptionally activated this pathway, while abatacept and, partly, tocilizumab were suppressive (**Figure 4E**). JAKi had additional activating effects on HDACs and the genes of glycolysis, and PPP. Tocilizumab treatment suppressed the genes of glycolysis pathway in CD4^+^ cells.

JAKi and abatacept treatment increased the transcription of *CCR7, CD27* and *IL23A*, activating the memory phenotype of the T cells (**Figure 4F**). The immune response genes *SOCS2*, *EGR1*, *IL23A*, *CD3G, NUP210* and *SCIMP* were transcriptionally affected by treatment with JAKi, methotrexate and abatacept, however the effects were frequently opposing. Among the insulin-responsive gene subset, JAKi and methotrexate suppressed transcription of chromatin remodeling genes. JAKi negatively regulated *KMT2A*, *PRKDC*, and *RIF1*, and activated *EZH2* and *ARID1A,* while methotrexate and abatacept suppressed *EZH2, ARID1A* and *PRKDC* (**Figure 4G**). The exhaustion receptor genes *PDCD1*, *TIGIT* were suppressed by methotrexate, *CTLA4* gene was suppressed by abatacept. In contrast, JAKi activated *PDCD1*. The IL6-receptor inhibitor tocilizumab did not affect the transcription of the insulin-responsive genes or the T cell exhaustion receptors suggesting that these effects are independent of the IL6 pathway in CD4^+^ cells.

Anti-rheumatic drugs had diverse effect on specific markers of the metabolic active synovial BIRC5^hi^CD4^+^ clusters (**Supplementary Figure S2B**). Methotrexate suppressed Tph markers PDCD1, CXCL13, ZNRF1, and had no significant effect on other BIRC5^hi^CD4^+^ clusters. Abatacept exhibited suppressive effect on the proliferating cell cluster. JAKi, abatacept and tocilizumab activated transcription of the *GNLY, GZMB, NKG7, KLRD1* genes specific for the Vdelta 1+2 and GLNY^+^CD4^+^ clusters. JAKi transcriptionally activated all metabolic active clusters, which was consequent to their insulin enhancing properties (6) and the effect on transcriptional profile of the metabolic pathway genes (**Figure 4E**).

Together, analysis of effect of anti-rheumatic drugs on transcriptome of CD4^+^ cells suggest that methotrexate, abatacept and JAKi only partly mitigated the insulin-responsive pathways i.e., chromatin remodeling and immune response with focus on control of T cell exhaustion and memory phenotype development. The genes of joint-specific clusters migrating to synovia were also differentially affected by anti-rheumatic treatment.

## Discussion

In this study, we link the insulin response of CD4^+^ cells to their pathogenic conversion in canonical autoimmune RA. Changes in the transcriptome induced by insulin are dependent on the genomic binding of survivin to H3K27ac. The regulatory elements of these genes adopt survivin binding based on insulin signaling and coordinate the pathways of T cell maturation and chromatin remodeling, thereby strengthening the link between survivin activity within regulatory chromatin (50) and the glucose metabolism control in CD4^+^ cells reported recently (5,6). Additionally, a subset of these genes defines the pathogenic insulin-responsive gene signature identifiable in CD4^+^ cells of classical insulin-resistant T2D and in the blood, synovial fluid, and synovial tissue of RA patients.

The study demonstrated that survivin-dependent transcriptional control of the insulin dependent genes was realized in parity with deposition of the H3K27ac histone modification, reported to drive the aggressiveness of active and proliferating T cells (8). Insulin responsiveness in RA survivin/*BIRC5^hi^* CD4^+^ cells activates histone acetylation enzymes, upregulating the glycolysis and the pentose phosphate pathway, the primary mechanisms of survivin-dependent metabolic adaptation employed by CD4^+^ T cells in RA (5). We simulated an insulin-sufficient scenario by insulin stimulation of healthy CD4^+^ cells *ex vivo* or through continuous linear regression of plasma insulin levels to the transcriptional profile of CD4^+^ cells in RA patients. Sufficiency of insulin reduced the transcription of the genes controlling histone methylation such as *KMT2A* and *EZH2*, proteins that promote development of a memory T cell phenotype (20,51). In agreement with this, our recent reports have also shown that survivin binds the catalytic center of *EZH2*, changing the histone methylation landscape (18,50), paralleling survivin’s role in insulin signaling. Insulin stimulation permits activation of memory markers *CD27, S1PR1, HOBIT/ZNF683* and suppress exhaustion markers *PDCD1* and *CTLA4*. Inhibition of HDAC has an inverse relationship to the downregulating effects of insulin, revealing the synergistic and protective effect of insulin and histone deacetylation in RA. Chromatin binding of survivin straddles the molecular events of increased metabolic activity in CD4^+^ T cell clusters and insulin-dependent histone acetylation in facilitating a memory T cell phenotype. In a larger context, our study adds value to the emerging links between metabolism and epigenetic remodeling in autoimmunity (1,52–54).

Histone acetylation figured prominently as a differentially modified epigenomic region in RA synovial fibroblasts of RA compared to non-RA (55). The study reported the cluster-wise distribution of epigenomic landscape profiling DNA methylation, several histone modifications, open chromatin regions, gene expression within functional genomic regulatory elements (55). Here, we connect histone acetylation to insulin signaling and metabolic activity of T cell clusters in the RA synovial tissue, synovial fluid, and blood, analyzing it on a single-cell level. In synovial tissue of RA, we demonstrated a diversity of the survivin/*BIRC5*-dependent profiles among the T cell clusters. The cluster of PD1^+^CD4^+^ T cells, cytotoxic CD4^+^GZMK^+^ and CD4^+^GNLY^+^ cells, the Vdelta cells and the proliferating cells reflected a metabolically active state in blood and showed enrichment in the synovial tissue in RA. Further, *BIRC5^hi^* T cells in blood of RA patients presented enrichment of these metabolically active T cells. Together, this suggests a direct trafficking mechanism of these cells into the inflamed joint.

Parallel to the effect of insulin and survivin on transcription detailed above, existing anti-rheumatic drugs methotrexate, JAKi and abatacept create a memory like phenotype through CD27 and *CCR7* upregulation and alleviate T cell exhaustion by suppressing C*TLA4, TIGIT and CD3G.* Further, JAKi and methotrexate suppress the transcription of genes that are responsive to insulin and regulated by histone acetylation such as *EZH2*, *PRKDC*, *ARID1A*, *NUP210*, exposing a targetable and easily detectable transcriptional signature based on insulin levels in the blood. Population cohort studies show that metformin affords blood glucose control through increased insulin sensitivity and relates to a 30% reduced risk of RA development (56,57). Analysing the transcriptome of healthy CD4^+^ cells, we found that anti-diabetic drug metformin also promoted a memory-like phenotype by activating *CD27* and *SELL* and suppressed cytotoxic *PRF1* and *GZMB,* highlighting that repurposing anti-diabetic drugs can be expedient in RA. On one hand, the changes in insulin-induced gene expression allow for optimization of RA treatment in active phase of the disease. On the other hand, our findings encourage new studies on the survivin-dependent insulin response as a reversible phase of RA to change the nature of aggressive PD1^+^, GNLY^+^, GZMK^+^ T cells aiming to prevent their entrance into joints.

In conclusion, our study presents a molecular epigenomic mechanism bridging the arthritogenic function of intracellular survivin in modulating insulin responsiveness and demarcating pathogenic T cell subclusters. In upcoming investigations, one needs to address in detail ability of antirheumatic drugs to modulate insulin sensitivity linking it to re-profiling of pathogenic T cells and RA disease regress.

## Figure legends

**Supplementary Figure S1.**
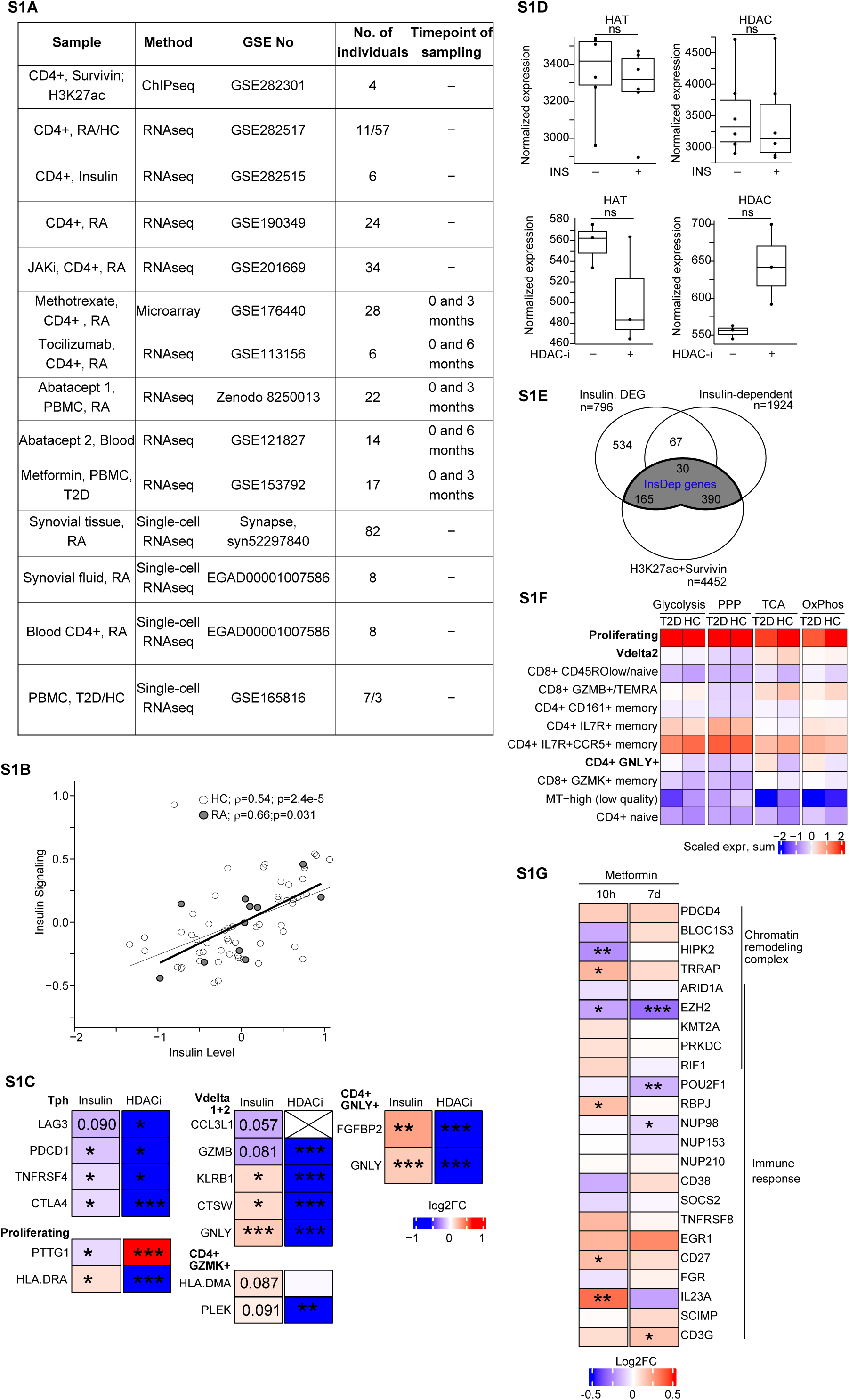
A. Table of patient material used in the analysis. B. Scatterplot showing spearman correlation and p-values between insulin levels in blood and summarized expression of insulin signaling genes (INSR, IRS1, IRS2) in patients with RA and healthy controls (HC). C. Heatmap of expression difference (by RNA-seq log2FC) of the markers of the synovial BIRC5^Hi^CD4^+^ clusters stimulated with insulin or treated with HDAC inhibitor romidepsin. Asterisks indicate Wilcoxon paired p-values, *<0.05, ** < 0.01, *** < 0.001 D. Boxplot showing normalized expression (by RNAseq) of HAT and HDAC enzymes in CD4^+^ cells stimulated with insulin (top) or treated with HDAC inhibitor romidepsin (bottom). Asterisks indicate Wilcoxon paired p-values, *<0.05, ** < 0.01, *** < 0.001 E. Venn diagram of insulin-dependent genes (blue, related to Figure 2H) connected to *cis*-RE containing survivin and H3K27ac deposition in chromatin of CD4^+^ T cells. F. Heatmap of expression sum of metabolic pathway enzymes in T cells of T2D patients and healthy controls (GSE165816). BIRC5^Hi^CD4^+^ cell clusters identified in synovia tissue of RA patients in Figure 1 are marked in bold. G. Heatmap of expression difference (by RNA-seq log2FC) of the insulin-dependent genes of the enriched pathways in CD4^+^ cells of healthy individuals treated with metformin short-term (10h) and long-term (7 days). Asterisks indicate Wilcoxon unpaired p-values, *<0.05, ** < 0.01, *** < 0.0001

**Supplementary Figure S2.**
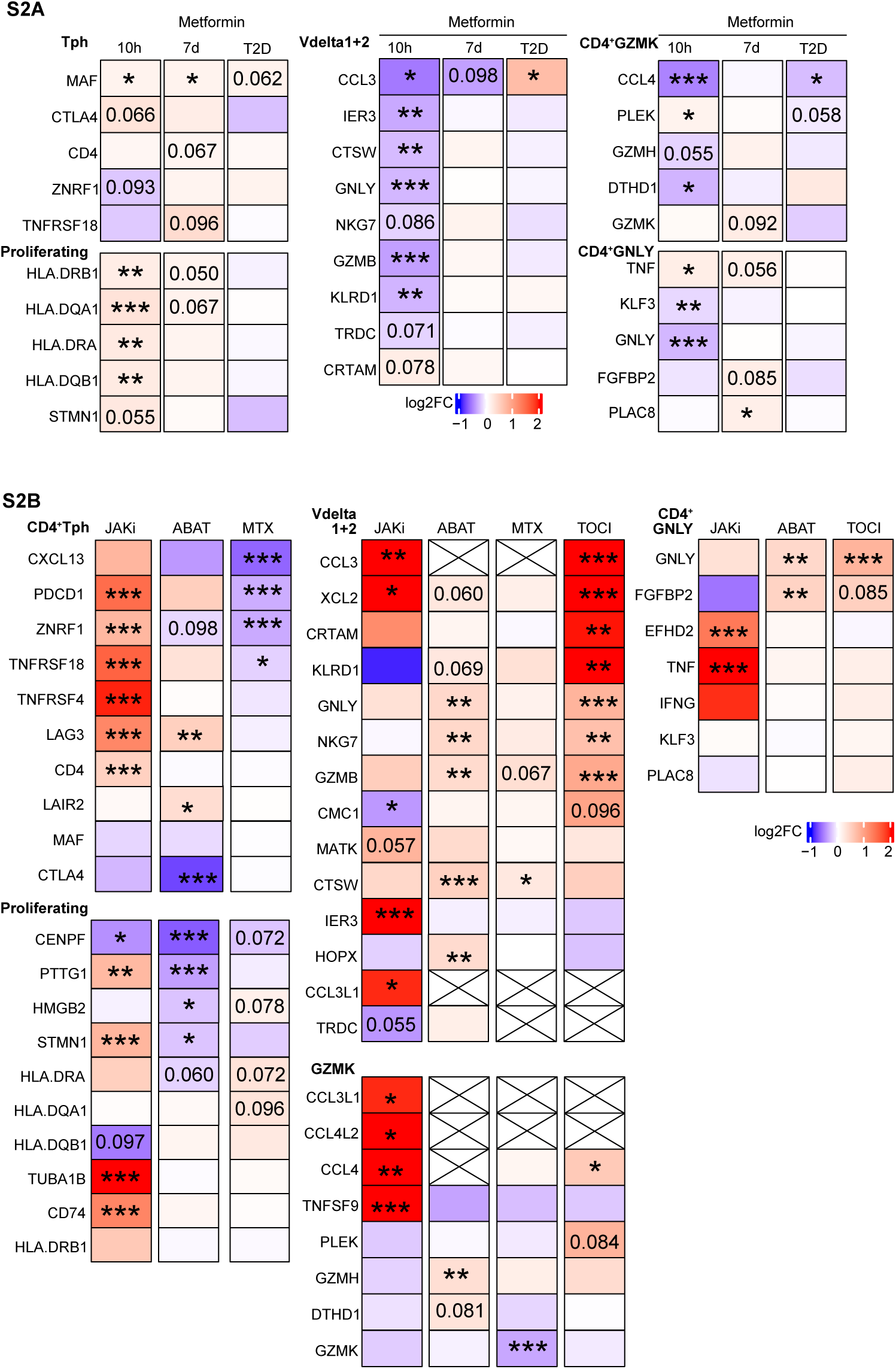
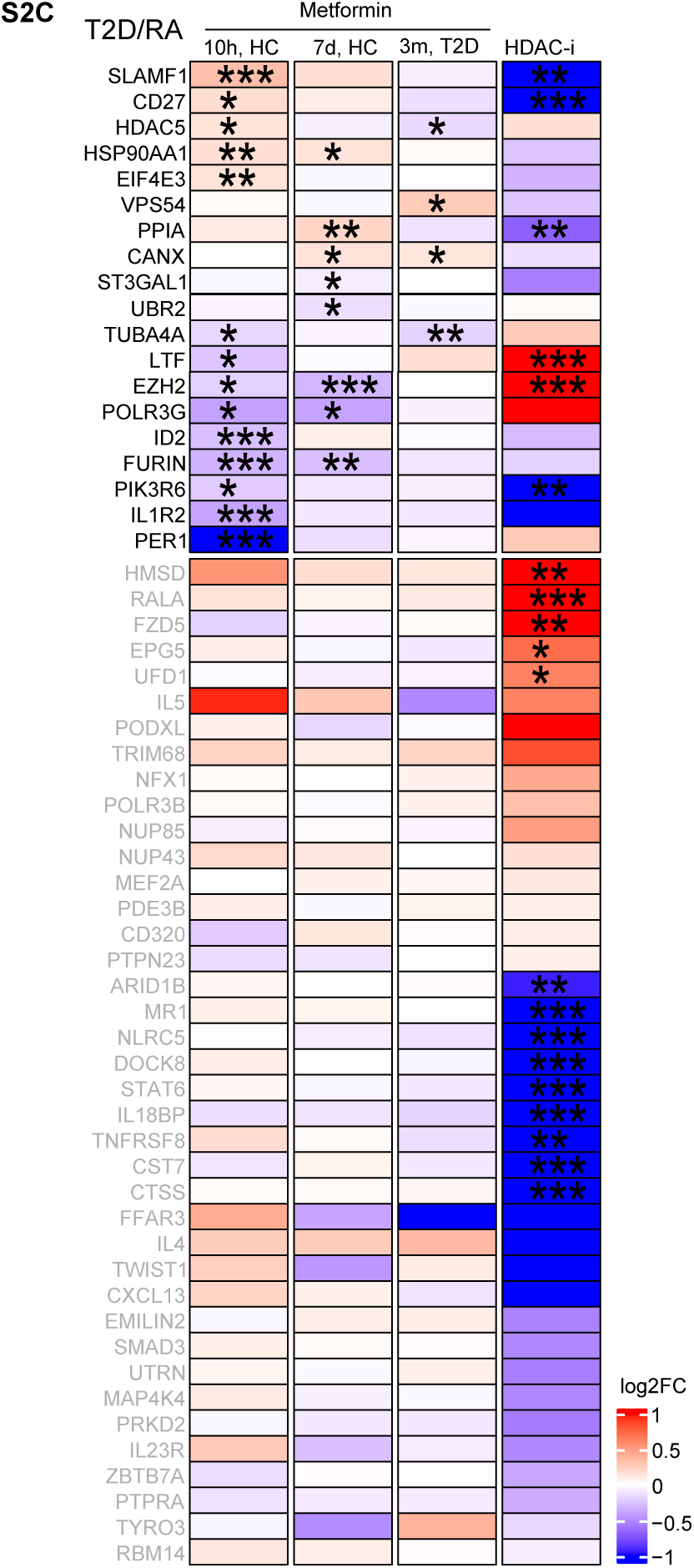
A. Heatmap of expression difference (by RNA-seq log2 fold change, FC) of the markers of the synovial BIRC5^Hi^CD4^+^ clusters of healthy individuals treated with metformin short-term (10h), long-term (7 days), and in T2D patients (3 months). Asterisks indicate Wilcoxon unpaired p-values, *<0.05, ** < 0.01, *** < 0.0001 B. Heatmap of expression difference (by RNA-seq log2FC) of the markers of the synovial BIRC5^Hi^CD4^+^ cell clusters of RA patients treated with JAKi (35 treated, 5 untreated), abatacept (ABA, n=22, paired), methotrexate (MTX, n=28, paired), tocilizumab (TOCI, n=6, paired). Wilcoxon p-values are shown. *< 0.05, **< 0.01, ***< 0.001 C. Heatmap of mean expression difference (by RNA-seq log2FC) of common T2D/RA genes (shown in Figure 4A) in CD4^+^ cells of healthy individuals treated with metformin short-term (10h), long-term (7 days), in T2D (3 months), and in CD4^+^ cells treated with HDAC inhibitor romidepsin. Asterisks indicate Wilcoxon paired test p-values, *< 0.05, **< 0.01, ***< 0.001

